## Supplemental Materials for "Selection-driven trait loss in independently evolved cavefish populations"

*Supplementary Methods: Inferring selection*

**hapFLK**

We conducted additional genome scans for selection using hapFLK (Fariello et al. 2013). The hapFLK statistic provides a powerful approach to detect regions of the genome under selection by testing for differentiation among populations in haplotype cluster frequencies exceeding what is expected by neutral evolution. We included a single *Astyanax aeneus* to serve as an outgroup, and we analyzed the Lineage 1 and Lineage 2 populations in separate hapFLK runs.

**SIFT/VEP**

We identified nonsynonymous coding variants across all populations and predicted the consequence of each variant on protein function using computational analysis with the SIFT (sorting intolerant from tolerant) algorithm (Ng and Henikoff 2003) and the Ensembl Variant Effect Predictor (VEP) software suite using Ensembl v100 annotations (McLaren et al. 2016). SIFT uses sequence homology and data on the physical properties of a given protein to predict whether an amino acid substitution will be tolerated or deleterious. VEP performs annotation and analysis of genomic variants to predict impact on the protein sequence (i.e., modifier, low, moderate, or high) (Table S4).

*Supplementary Results: Inferring Recent Demographic History*

Population genomic approaches used to detect signatures of selection commonly rely on demographic parameters inferred from the site frequency spectrum (SFS). We used a non-parametric method implemented in Stairway Plot 2 (Liu and Fu 2020) to estimate past changes in population size that may influence our ability to detect and classify regions of the genome that have experienced selective sweeps. We used previously generated unfolded SFS for Pachón cave, Tinaja cave, Molino cave, Río Choy, and Rascón populations (Herman et al. 2018). The white long fin tetra was used to infer the ancestral allele and polarize the data, which consisted of 500Mb of sequence (including invariant sites) for each population. The SFS for each population was provided to Stairway Plot 2 and singletons were masked. We used a generation time of 1 year and a mutation rate of 3.5e-9 estimated from cichlids (Malinsky et al. 2018).

Recently, (Herman et al. 2018) conducted demographic modeling in ∂a∂i (Gutenkunst et al. 2010) using 2D unfolded SFS to estimate present day effective population sizes and the timing of cave-cave, cave-surface and surface-surface (Lineage 1 and Lineage 2) population splits and timing of secondary contact (and admixture) between populations. The present analysis used the 1D unfolded SFS for each population to provide estimates of population size changes over time, as well as bottleneck and expansion events in the recent past. Cave populations were expected to have experienced marked bottlenecks corresponding to the time that the surface population invaded caves. Previous demographic modeling with ∂a∂i indicated that surface and cave populations from both Lineage 1 and Lineage 2 split from one another at remarkably similar times (~160,000 generations ago for both lineages), and that the split between the two surface lineages occurred around 260,000 generations ago. Previous analyses also supported a model of secondary contact with ongoing gene flow between cave and surface populations (Herman et al. 2018). We note that Stairway Plot 2 does not account for ongoing gene flow between populations, so estimates of effective population sizes may be inflated compared to previously published estimates from ∂a∂i (Herman et al. 2018).

Our estimates of past population size changes with Stairway Plot 2 indicated consistent differences between cave populations (i.e., Pachón, Tinaja, Molino) and surface populations (i.e., Rascón and Río Choy). Most notably, cave populations underwent a bottleneck corresponding to previously estimated timing of cave-surface population splits (i.e., invasion of caves by surface ancestors) between 100-200k generations ago (also supported by ∂a∂i analyses from (Herman et al. 2018)). Our stairway Plot 2 analysis revealed that both Lineage 2 (Rascón) and Lineage 1 (Río Choy) surface populations appear to have experienced a more ancient bottleneck event at approximately 800k generations ago, potentially corresponding to their migration into Northern Mexico, followed by subsequent population expansions.

*Supplementary Results: Determining the origins of Cueva del Río Subterráneo*

Recent introgression events can have a large impact on species tree inference. We recently found evidence that Chica and Caballo Moro caves contain surface-cave hybrids with 15-25% of their genomes derived from surface fish ancestry, on average ((Moran et al. 2022), Medley et al. in review). Previous studies have suggested that Subterráneo cavefish may also experience a substantial amount of admixture with the local surface population. Preliminary phylogenomic analyses indicated that Subterráneo grouped with surface populations rather than with other cave populations, and therefore may represent a unique evolutionary origin of cave adaptation distinct from the El Abra and Guatemala cave lineages. However, this pattern might also be driven by substantial ongoing gene flow with the local Micos River surface population.

Surface water floods into Subterráneo cave during the wet season, bringing *A. mexicanus* surface fish into the cave. Unlike most other cave populations (with the exception of Chica and Caballo Moro hybrid populations), cavefish within Subterráneo have been shown to have eyes present (although reduced in size compared to surface fish) and pigmentation ((Wilkens and Burns 1972) (Elliott 2018)). Surface fish have also been documented in Subterráneo cave during the wet season after high water. Although previous phylogenetic analyses have found support for Subterráneo cave as a close relative to the Lineage 1 cavefish in the Guatemala region, this cave occurs within El Abra limestone and is geographically closer to the El Abra caves compared to the Guatemala caves. Ongoing hybridization with Lineage 1 surface fish (i.e., from Arroyo La Pagua that floods into the cave, (Elliott 2018)) could be masking that this cave population originated from the same Lineage 2 surface fish stock that populated the El Abra caves (first proposed by (Coghill et al. 2014)).

To explore this hypothesis, we conducted formal tests for introgression between Subterráneo cavefish, a Lineage 1/Guatemala region cavefish population (Escondido), two Lineage 2/El Abra region cavefish population (Pachón and Tinaja, representing caves at the northern and southern extent of the El Abra cave region), a Lineage 1 surface fish population (Mante), and a Lineage 2 surface fish population (Rascón). Mante was included as the new lineage surface fish population in these analyses rather than Micos due to sample size (Micos: n=1; Mante: n=10). Two *Astyanax A. nicaraguensis* samples served as an outgroup comparison.

We conducted formal tests for introgression with Treemix and using D and f4 statistics. We first used Treemix v1.13 (Pickrell and Pritchard 2012) to visualize migration events and confirm phylogenetic relationships between the Subterráneo population and the two non-admixed cave populations, the two surface populations, and the outgroup. Treemix builds a bifurcating tree to represent population splits and also incorporates migration events, which are represented as “edges,” connecting population branches. For this analysis, we used biallelic SNPs thinned to 1kb apart. We supplied the resulting set of 1,148,321 SNPs to Treemix, rooted with *A. nicguensis*, and estimated the covariance matrix between populations using blocks of 500 SNPs. Sample Rascón _6 was excluded from this analysis because ADMIXTURE indicated that it was likely an early generation hybrid. We first built the maximum likelihood tree (zero migration events) and then ran Treemix sequentially with one through six migration events. We calculated the variance explained by each model (zero through six migration events) using the R script treemixVarianceExplained.R (Card 2015).

We used Dsuite v0.4 ^55^ to conduct formal tests for introgression between Subterráneo cavefish and new lineage and old lineage surface and cave populations. This allowed us to further test the hypothesis that Subterráneo represents a hybrid population resulting from admixture between a Lineage 2/El Abra region cave population and a Lineage 1 surface population. If gene flow has occurred between Subterráneo cavefish and the local Lineage 1 surface population, we predict an excess of shared derived alleles between Mante and Subterráneo. This analysis may also provide insight into which lineage of surface stock founded the Subterráneo cave population. If Subterráneo cave was initially populated by Lineage 2 surface fish, we would expect Subterráneo cavefish to share more derived alleles with Lineage 2 cave populations compared to Lineage 1 cave populations. However, we note that recent, ongoing introgression with the Lineage 1 surface fish may artificially inflate the number of shared derived alleles between Subterráneo and Lineage 1 cavefish.

We supplied the same set of 1,148,321 thinned biallelic SNPs used in the Treemix analysis to Dsuite and specified *A. nicaraguensis* as the outgroup. We again excluded the one sample from Rascón with apparent hybrid ancestry. We used the Dsuite program Dtrios to calculate Patterson’s D statistic for all possible trios of populations using the ABBA-BABA test ^56^. The ABBA-BABA test quantifies whether allele frequencies follow those expected between three lineages (e.g., sister species P1 and P2, and a third closely related species, P3) under expectations for incomplete lineage sorting (ILS). Observing a greater proportion of shared derived alleles between P1 and P3 but not P2 or between P2 and P3 but not P1 than what would be expected by chance (i.e., ILS) indicates introgression. Dsuite requires a fourth population, P4, to serve as an outgroup and determine which alleles are ancestral versus derived. Ancestral alleles are labeled as “A” and derived alleles are labeled as “B”. ABBA sites are those where P2 and P3 share a derived allele, and ABAB sites are those where P2 and P4 share a derived allele. The D statistic is calculated as the difference in the number of ABBA and BABA sites relative to the total number of sites examined. Dsuite uses jackknifing of the null hypothesis that no introgression has occurred (D statistic = 0) to calculate a p-value for each possible trio of populations.

Dsuite also calculates the admixture fraction, or f4-ratio, which represents the covariance of allele frequency differences between P1 and P2 and between P3 and P4. If no introgression has occurred since P1 and P2 split from P3 and P4, then f4 = 0. If the f4 statistic is positive, this suggests a discordant tree topology indicative of introgression.

We then quantified introgression across the genome in Subterráneo cavefish using Hidden Markov Model (HMM) and fine-scale SNP mapping approaches to calculate ancestry proportions globally (i.e., genome-wide averages) and locally (i.e., at each site along each of the 25 chromosomes). We implemented a HMM-based approach in Loter to infer genome-wide local ancestry in the Subterráneo individuals. Mante served as the parental surface population for the initial training stage of the HMM, as we only had a sample size of one for the local Micos surface population and our phylogenetic analyses revealed that Mante surface fish are closely related to the Micos and Subterraneo populations. As preliminary analyses indicated that Subterráeno shared more derived alleles with Lineage 2/El Abra region cavefish compared to

Lineage 1/Guatemala region cavefish, we ran the analysis with Pachón as the proxy for the parental cave population (i.e., representing the genome of Subterráneo cavefish prior to onset of recent admixture with the local surface population). This analysis allowed us to estimate global ancestry proportions and mean minor and major parent tract lengths for each individual. Ancestry tract lengths were converted from base pairs to Morgans using the median genome-wide recombination rate 1.16 cM/Mb (0.0000000116 Morgan/bp) obtained from a previously published genetic map for *A. mexicanus* ^63^. We then estimated the number of generations since the onset of admixture (T_admix_) using the following equation:

T_admix_ = 1/(L_M_*p_B_)

where L_M_ is the mean ancestry tract length from the minor parent in Morgans and p_B_ is the proportion of the genome derived from the major parent (the probability of recombining) ^64–66^. This analysis indicated an estimated mean ± SE of 9,843 ± 600 generations since the onset of admixture.

*Supplementary Results: Population Structure*

ADMIXTURE analysis revealed fine-scale patterns of genetic differentiation among populations. Of note, Japones and Yerbaniz cave populations grouped together, separate from the majority of the other El Abra cave populations (i.e., Jos, Montecillos, Palma Seca, Sabinos, Tigre, and Tinaja) which mirrors their geography, as they are less than 2 km from each other, possibly interconnected (Mitchell et al. 1977, Elliott 2018), and documented to share a genetic basis of the brown mutation (Gross et al. 2009). Pachón was also categorized as a distinct genetic group from the other El Abra caves. Chica and Toro, the two southernmost El Abra caves that are geographically very close to one another, were also grouped in this analysis. Admixed ancestry was evident in Toro, in agreement with previous work suggesting this cave contains recent hybrids between cave and surface fish (Panaram and Borowsky 2005). Admixed ancestry was not evident for the Chica cave population at K=11, but for some of the lower values of K (e.g., K=7) Chica individuals showed admixed ancestry with Río Choy surface (i.e., a proxy for Lineage 1 surface fish) and El Abra cave parental populations. This agrees with recent detailed analyses into the history of admixture in Chica cave (Moran et al. 2022).

PCA on SNPs revealed genetic clustering of samples into eight clades, most of which contained only surface or only cave populations (Fig. S4). PC1 accounted for 20.5% of the total variance and separated Lineage 1 from Lineage 2 populations. PC2 accounted for 13.2% of the total variance and separated the cave from the surface populations. Subterráneo cave was the only population that did not group with like ecotypes, instead clustering with several surface populations, including samples from Micos river. A tributary of the Micos surface population, Arroyo La Pagua, floods into Subterráneo cave during high water in the wet season, and it is common to observe surface fish in this cave (Elliott 2018). Thus, gene flow into Subterráneo cave from the surface could explain this pattern.

Notably, the Arroyo cave samples were split across two clusters, with some individuals grouping with old lineage cave populations, as expected from the geographic location of this cave, and other individuals clustering with new lineage surface populations. This variation within samples from Arroyo is likely due to ongoing hybridization between this cave population and the local surface population. When fish were collected from Arroyo cave in 2019, putative hybrids were observed with some asymmetry in eye loss and variation in body pigmentation and eye degeneration (P.O.-G., pers. obs.). Some surface fish, including cichlids, were also encountered in the cave pools, indicating a connection with the local surface waters, and some *Astyanax* cavefish showed parasites clearly translocated from the surface fish (P.O.-G., pers. obs.).

As expected given their smaller populations sizes and historical bottleneck events, cavefish populations tend to have much lower nucleotide diversity (pi) compared to surface fish populations (Table S19, Figure S17). The genome-wide average for pi varied by an order of magnitude among all *A. mexicanus* populations, ranging from 0.00041 in Escondido cave to 0.0039 in the Mante surface population (Table S19, Figure S17). Mean D_XY_ ranged from 0.00045 between Jineo and Molino caves (both Lineage 1 and in the Guatemala region) to an order of magnitude larger (0.0052) between Mante (Lineage 1) and Peroles (Lineage 2) surface populations (Table S19, Figure S18A). These patterns are consistent with hypotheses surrounding the time since divergence and degree of gene flow between populations. Pairwise F_ST_ values (Table S19, Figure S18B) largely reflect the high degree of variation in pi among populations rather than variation divergence, and thus, pairwise F_ST_ is not a reliable metric to quantify the divergence between populations in this system (Charlesworth 1998, Herman et al. 2018).

*Supplementary Results: Cueva del Río Subterráneo Admixture Analyses*

While our phylogenetic analyses suggested that Subterráneo cave may represent a third independent origin of cave adaptation in *Astyanax* (Figures 1C, S1-S3), we found strong support for an alternative hypothesis. Specifically, our analyses indicated that recent admixture between the Subterráneo cave and the Micos surface population, a tributary of which (i.e., Arroyo La Pagua), floods into Subterráneo cave during high water in the wet season, has caused Subterráneo cavefish to group phylogenetically with surface fish populations rather than with other cavefish populations.

We inferred historical and contemporary migration events between Subterráneo cavefish, two Lineage 2 cave populations in the El Abra region (Pachón and Tinaja, representing caves at the northern and southern extent, respectively, of the El Abra cave region), a Lineage 2 surface population (Rascón), a Lineage 1 surface population (Mante), and an outgroup (*Astyanax nicaraguensis*). This analysis indicated historic gene flow events between Subterráneo and the El Abra caves (Pachón and Tinaja) and recent gene flow between Subterráneo and Mante, the Lineage 1 surface fish.

Formal tests for introgression using D and f4 statistics found significant support for introgression between all populations examined. For a given population trio, we observed that Subterráneo shares a higher percentage of derived alleles with the Lineage 1 cavefish population (Escondido: 33%) compared to the Lineage 2 cavefish populations (Pachón: 26%; Tinaja: 23%) (Table S2). This could be interpreted to suggest that Subterráneo cavefish originated from new lineage surface stock. However, ongoing hybridization with the local Lineage 1 Micos surface fish may artificially inflate the number of shared derived alleles present between Subterráneo cavefish and the Lineage 1 cavefish.

Fine-scale ancestry mapping using a HMM also indicated that Subterráneo individuals appear to have a highly admixed genome with ancestry from the Lineage 1 Micos surface population and Lineage 2/El Abra region caves (Figure S6), indicating that Subterráneo cave was originally populated by the same Lineage 2 surface stock that populated the El Abra caves. Notably, our analyses showed that the Subterráneo hybrid population is unique from other previously described hybrid populations in that the majority of ancestry is derived from surface fish. In contrast, hybrid populations in Chica and Caballo Moro caves were recently shown to have a majority of their genomes derived from cave ancestry (Moran et al. 2022, Medley et al. in review).

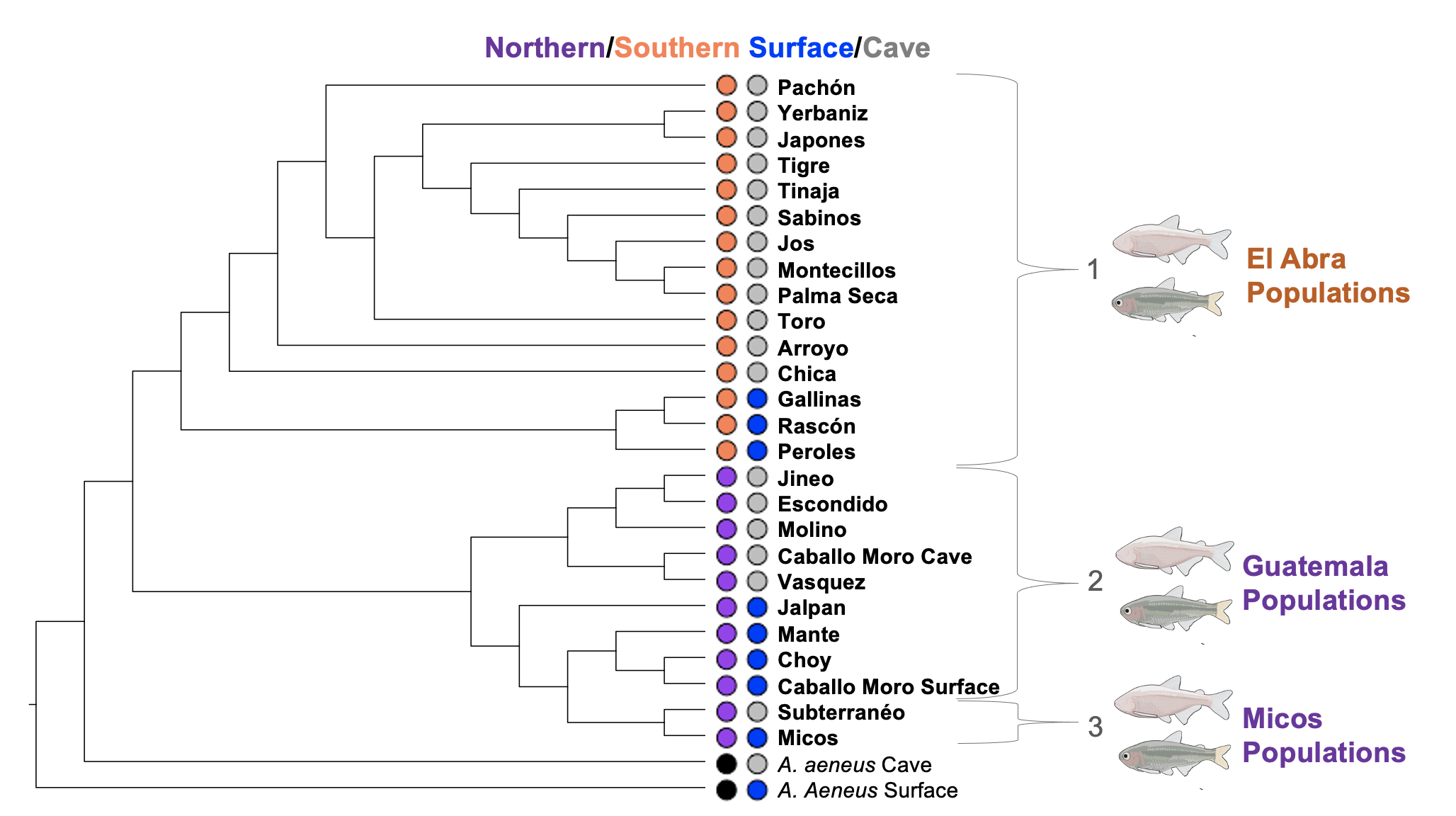

**Figure S1.** Phylogenomic signature of repeated evolution of cave adaptation in A. mexicanus. Shown is the maximum likelihood population tree built in Treemix using 680,021 SNPs shared across all populations sampled.

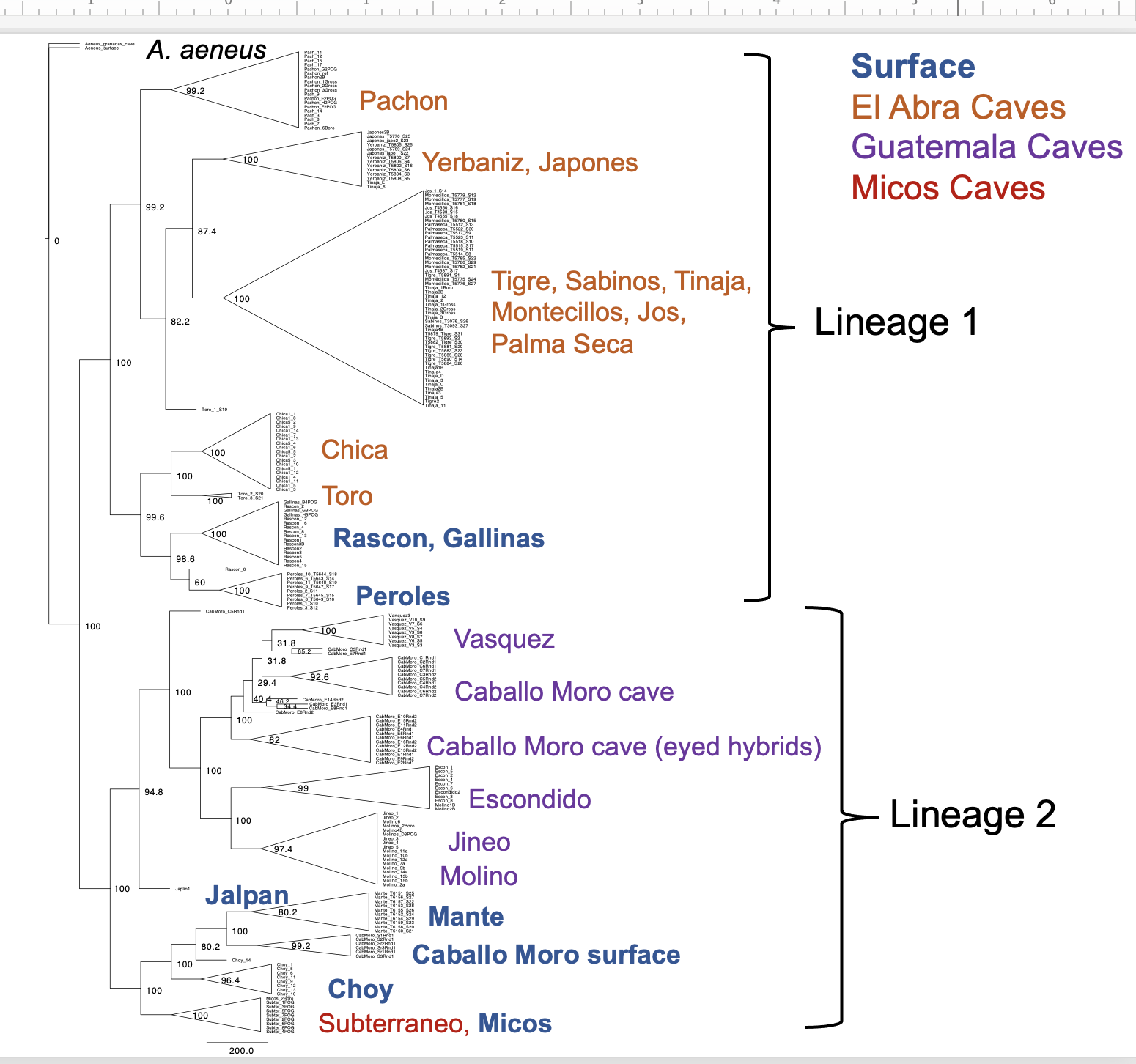

**Figure S2.** Full multi-species-coalescent tree with all individuals labeled, inferred using 1,121,282 SNPs in SVDQuartets**.** SVDquartets was run with a sampling of 500,000 random quartets and 500 standard bootstrap replicates specified to obtain bootstrap node support

values.

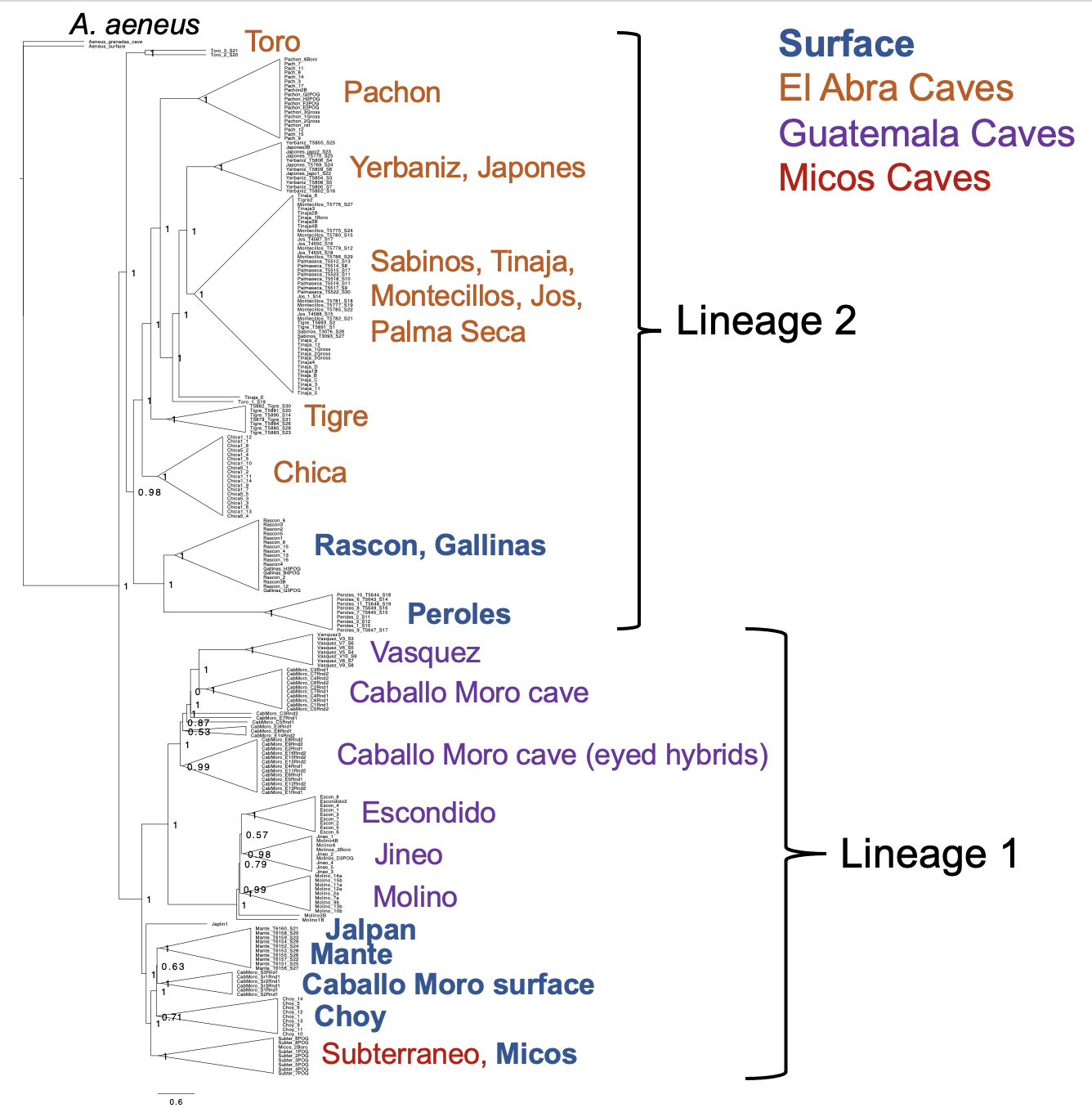

**Figure S3.** Species tree inferred using a coalescent gene tree-based approach (using 3,339 single copy orthologs covering 63,395,097 BPs total) in ASTRAL. Branch lengths are given in coalescent units and branch supports (node labels) are measured as local posterior probabilities. Final quartet score = 440574139629. Final normalized quartet score = 1.047.

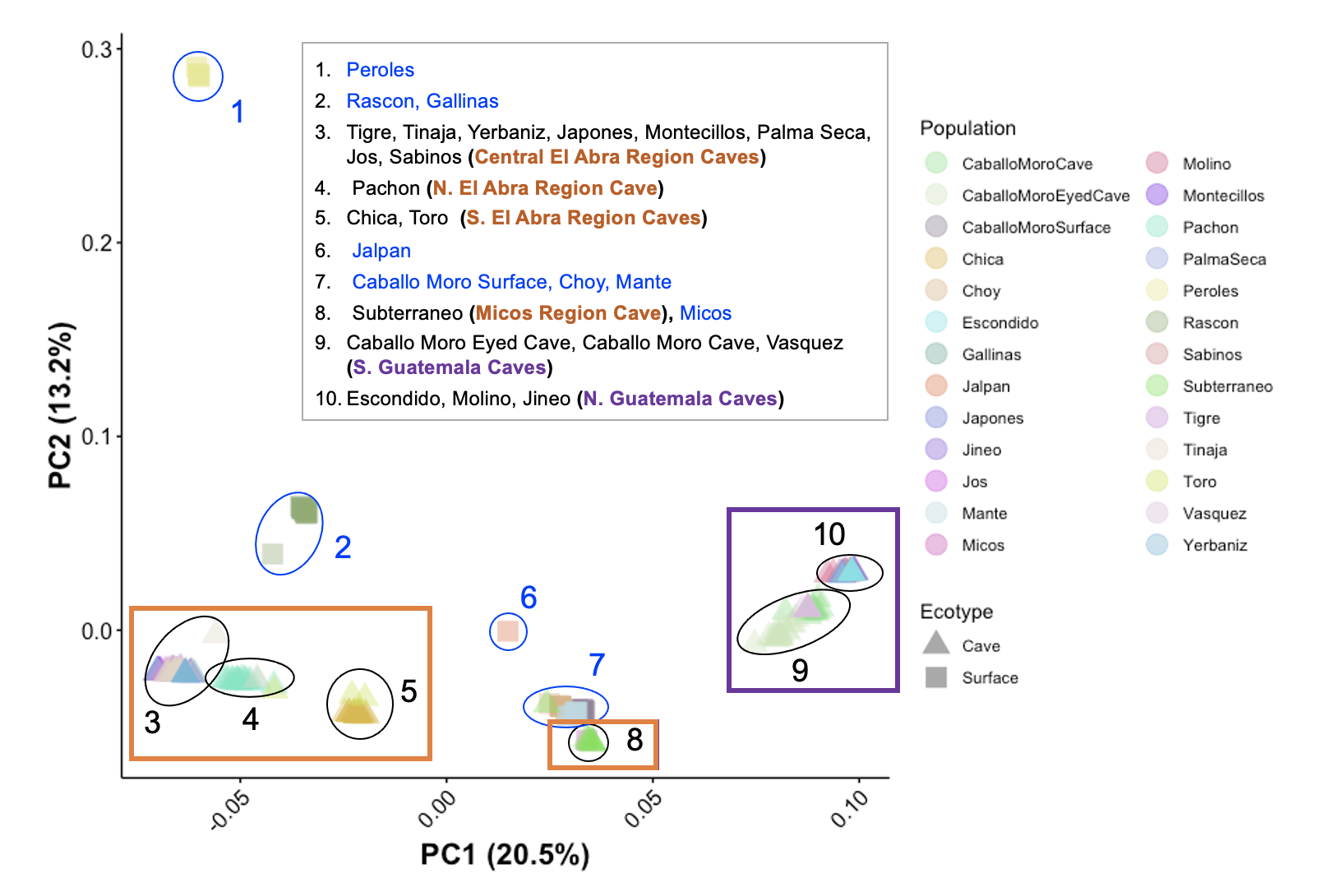

**Figure S4.** Principal Component (PC) analysis on 751,759 SNPs across eight surface and 18 cave populations of *Astyanax mexicanus* (see Table S1 for sample sizes)*.* Each population is assigned a unique color (see legend). For surface populations, the collection location is provided in blue text in the inset and individuals are represented with squares. For cave populations, the collection location is provided in black text in the inset and individuals are represented with triangles. Additionally, the Micos and Guatemala caves are shown within purple boxes (these two cave regions contain cavefish originating from the same lineage of surface stock). The El Abra caves are shown within an orange box (this cave region contains cavefish from a second independent lineage of surface stock). Clustered populations are grouped in clades numbered 1-10 (indicated with ellipses; also see inset) based on overlapping PC values.

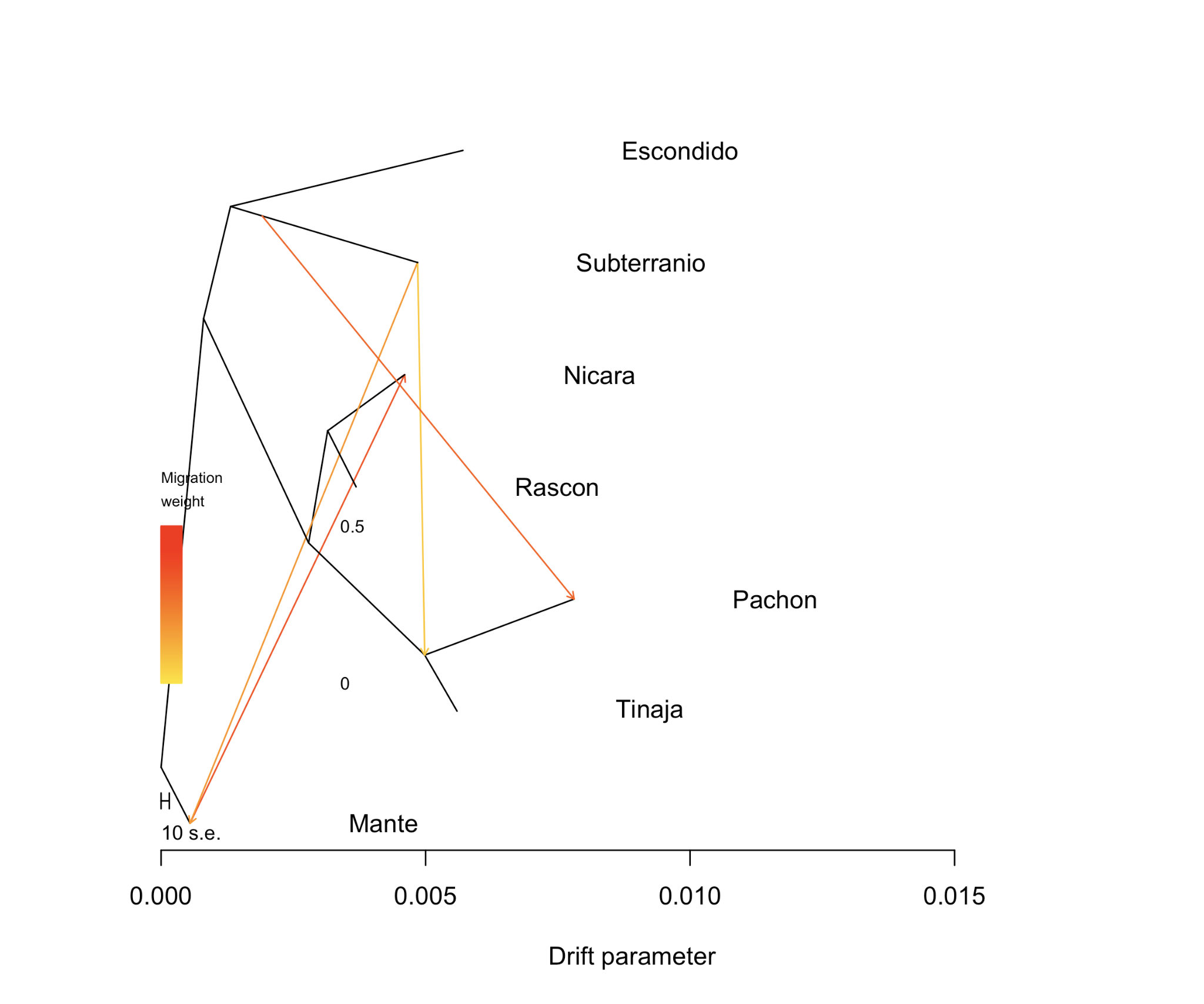

**Figure S5.** Treemix graph depicting the phylogenetic relationship and inferred migration events (m=4) between Subterráneo, two Lineage 2 cave populations in the El Abra region (Pachón and Tinaja, representing caves at the northern and southern extent, respectively, of the El Abra cave region), a Lineage 2 surface population (Rascón), a Lineage 1 surface population (Mante), and an outgroup (*Astyanax nicaraguensis*). Historic gene flow events are indicated between the El Abra caves (Pachón and Tinaja) with Subterráneo and recent gene flow is indicated between Subterráneo and Mante (representing Lineage 1 surface fish). Recent hybrids, as indicated by ADMIXTURE analysis, were removed from the data set prior to running additional analyses to detect introgression.

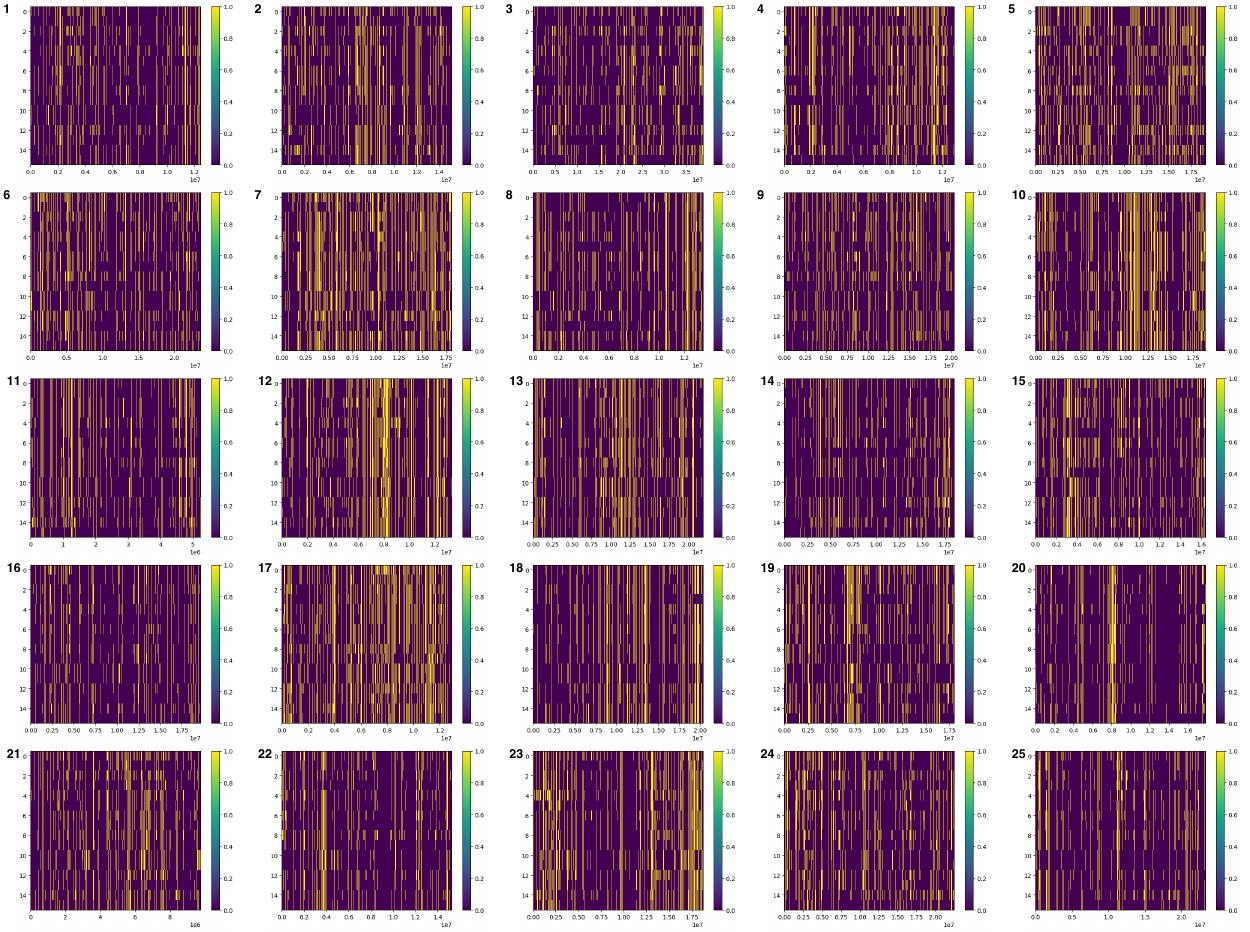

**Figure S6.** Visualization of local ancestry tracts in Subterráneo samples inferred using a Hidden Markov Model approach along each of the 25 chromosomes. Pachón and Mante were used as the cave and surface parental populations, respectively. Yellow represents cave ancestry and purple represents surface ancestry. The y axis shows haplotypes 0-15 corresponding to n = 8 diploid individuals. The x axis shows bp position along each chromosome.

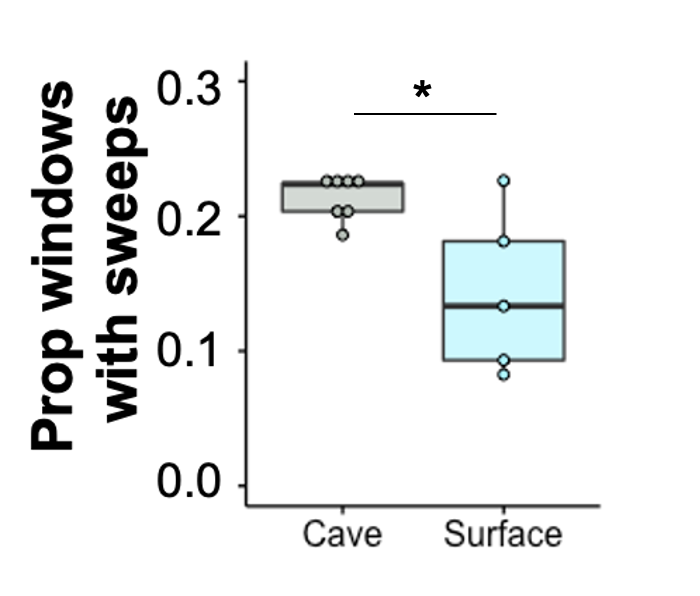

**Figure S7.** A significantly higher proportion of 5 kb genomic windows were predicted by diploS/HIC to contain a hard or soft selective sweep in cave (n=7) compared to surface (n=5) populations. * indicates p < 0.05. Mean ± SE proportion of 5 kb windows under selection in surface populations = 0.143 ± 0.027; mean ± SE proportion of 5 kb windows under selection in cave populations = 0.213 ± 0.006; Wilcoxon rank sum test: W = 29, p-value = 0.037.

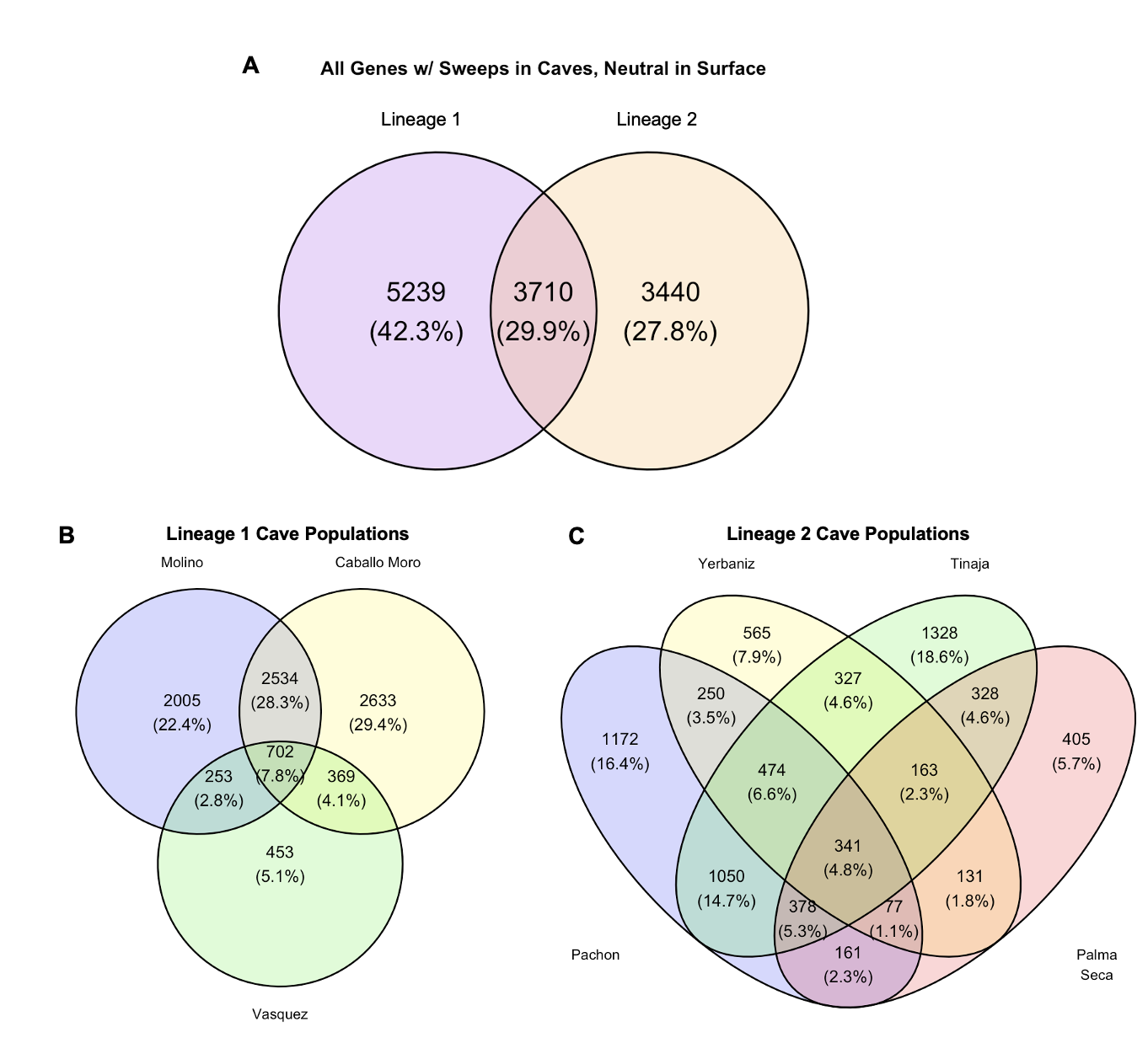

**Figure S8.** Venn diagrams showing overlap in number of genes with a sweep in a cave population and neutral evolution in a same-lineage surface population. (A) All Lineage 1 and Lineage 2 cave populations are grouped together. (B) Comparison among Lineage 1 cave populations. (C) Comparison among Lineage 2 cave populations.

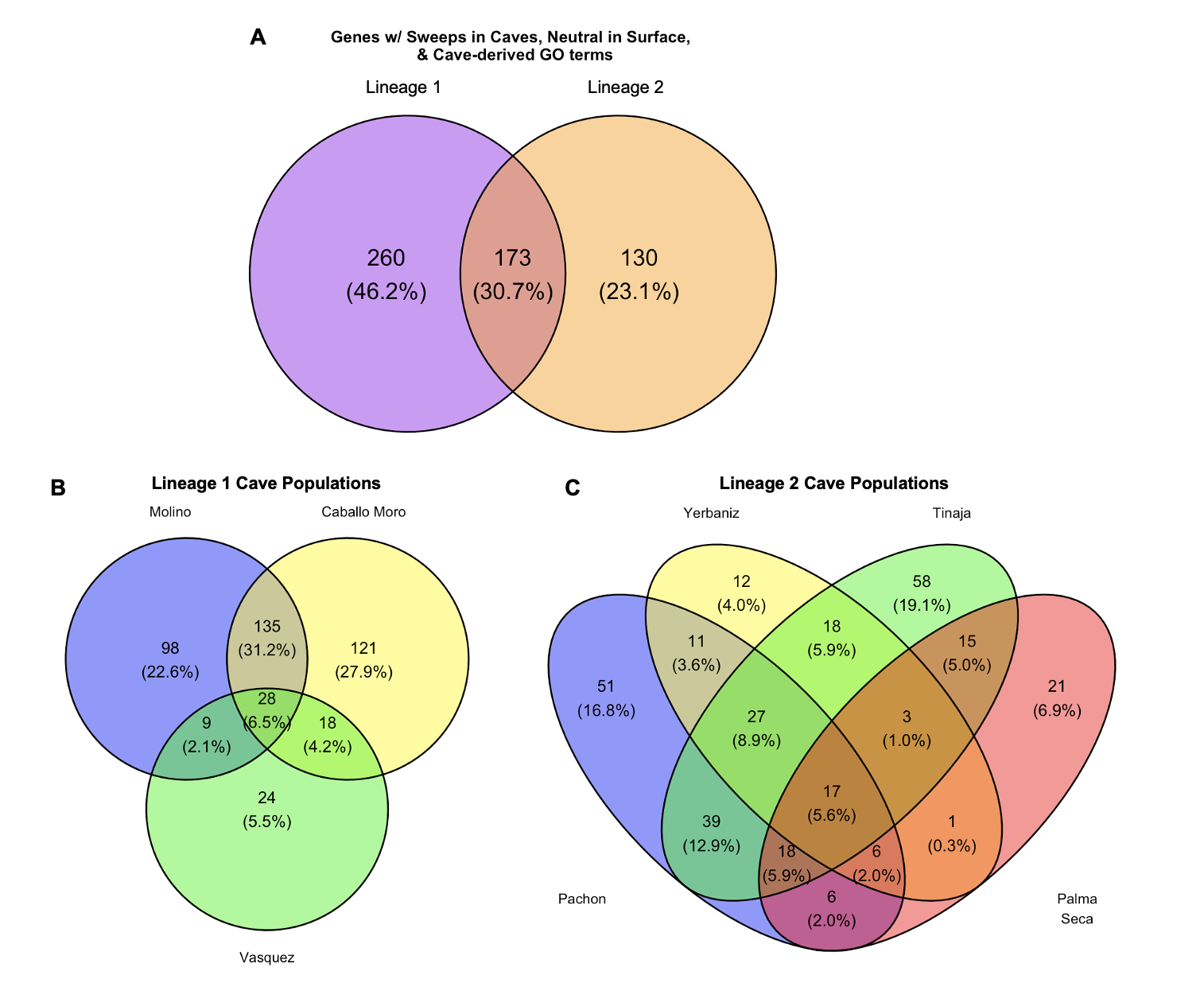

**Figure S9.** Venn diagrams showing overlap in number of genes with a sweep in a cave population, neutral evolution in a same-lineage surface populations, and relevant GO term associated with known cave-derived traits. (A) All Lineage 1 and Lineage 2 cave populations are grouped together. (B) Comparison among Lineage 1 cave populations. (C) Comparison among Lineage 2 cave populations.

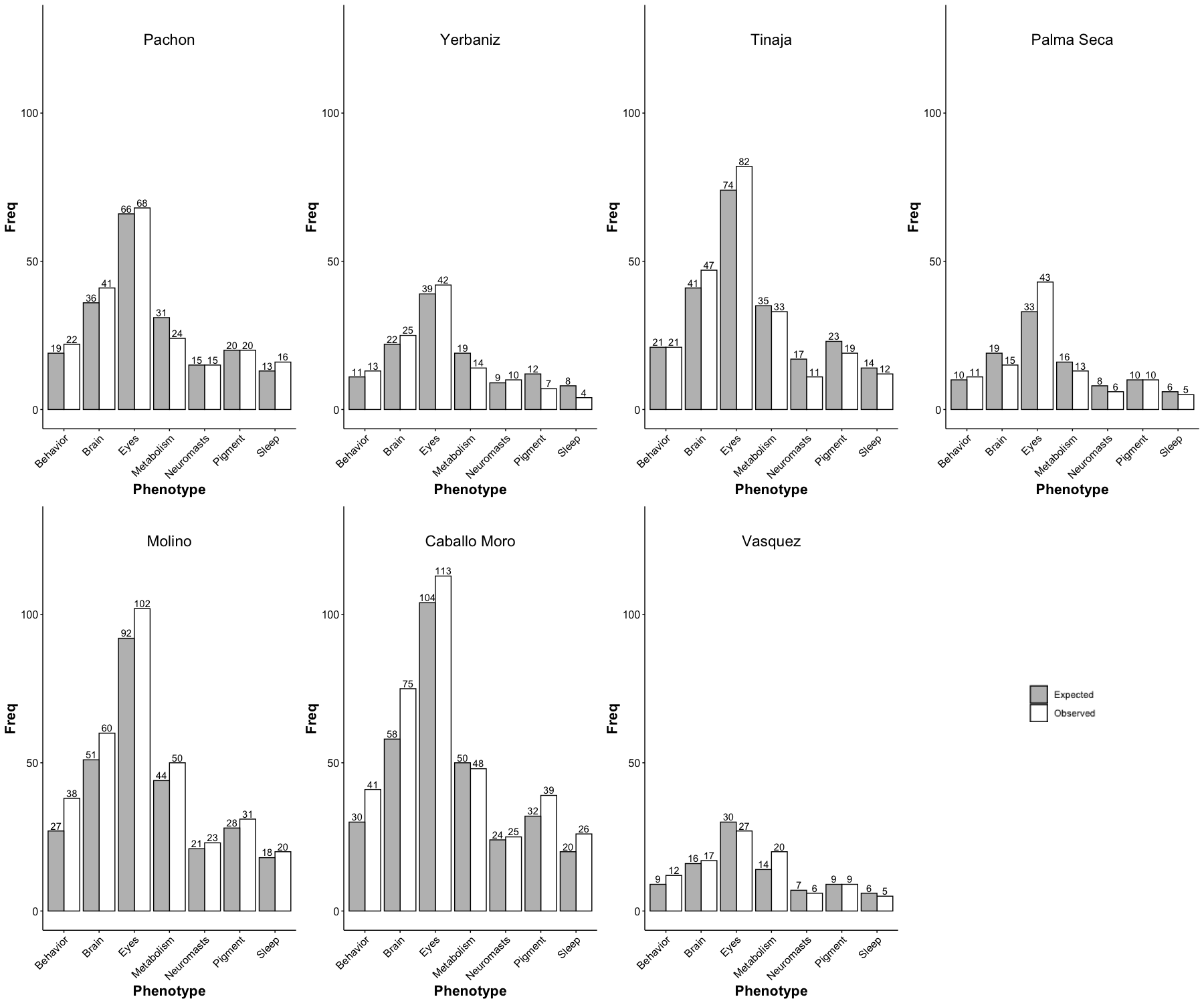

**Figure S10.** Number of expected and observed genes with GO terms matching a given phenotypic category associated with cave-derived traits in each of the seven cave populations examined. For each cave population, expected numbers for phenotypic categories were obtained by calculating the proportion of genes with GO terms in each phenotypic category in the entire surface fish genome annotation (26,698 gene total) and then multiplying by the total number of sweeps in each population. Fisher’s exact tests were performed for each phenotypic category within each population and no significant differences were found between the expected and observed counts. Note that pleiotropic genes (i.e., those with GO terms associated with 2 or more phenotypic categories) were included multiple times.

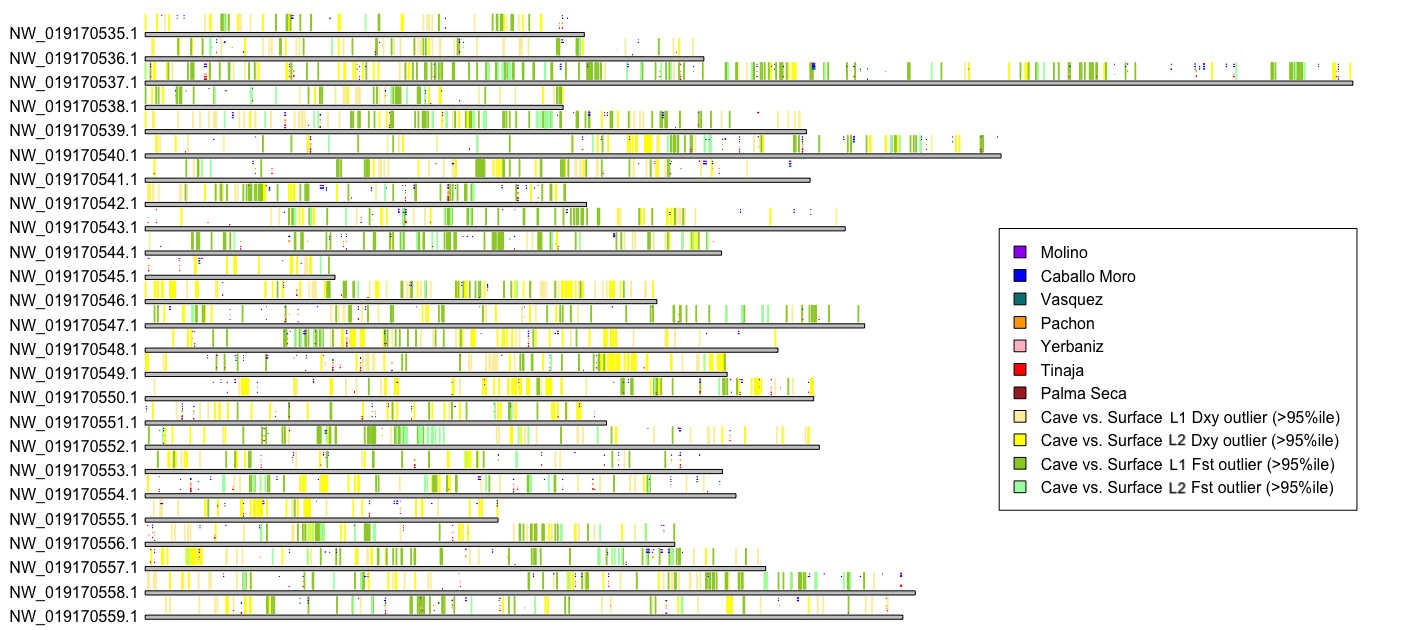

**Figure S11.** Locations of adaptive alleles (i.e., genes with selective sweeps and GO terms associated with cave-derived traits) across seven cave populations and outlier windows with exceptionally high divergence (Dxy and Fst values above the 95^th^ percentile, calculated in 50 kb windows) between cave and surface populations (i.e., comparison between all Lineage 1 cave individuals vs. all Lineage 1 surface individuals, and all Lineage 2 cave individuals vs. all Lineage 2 surface individuals).

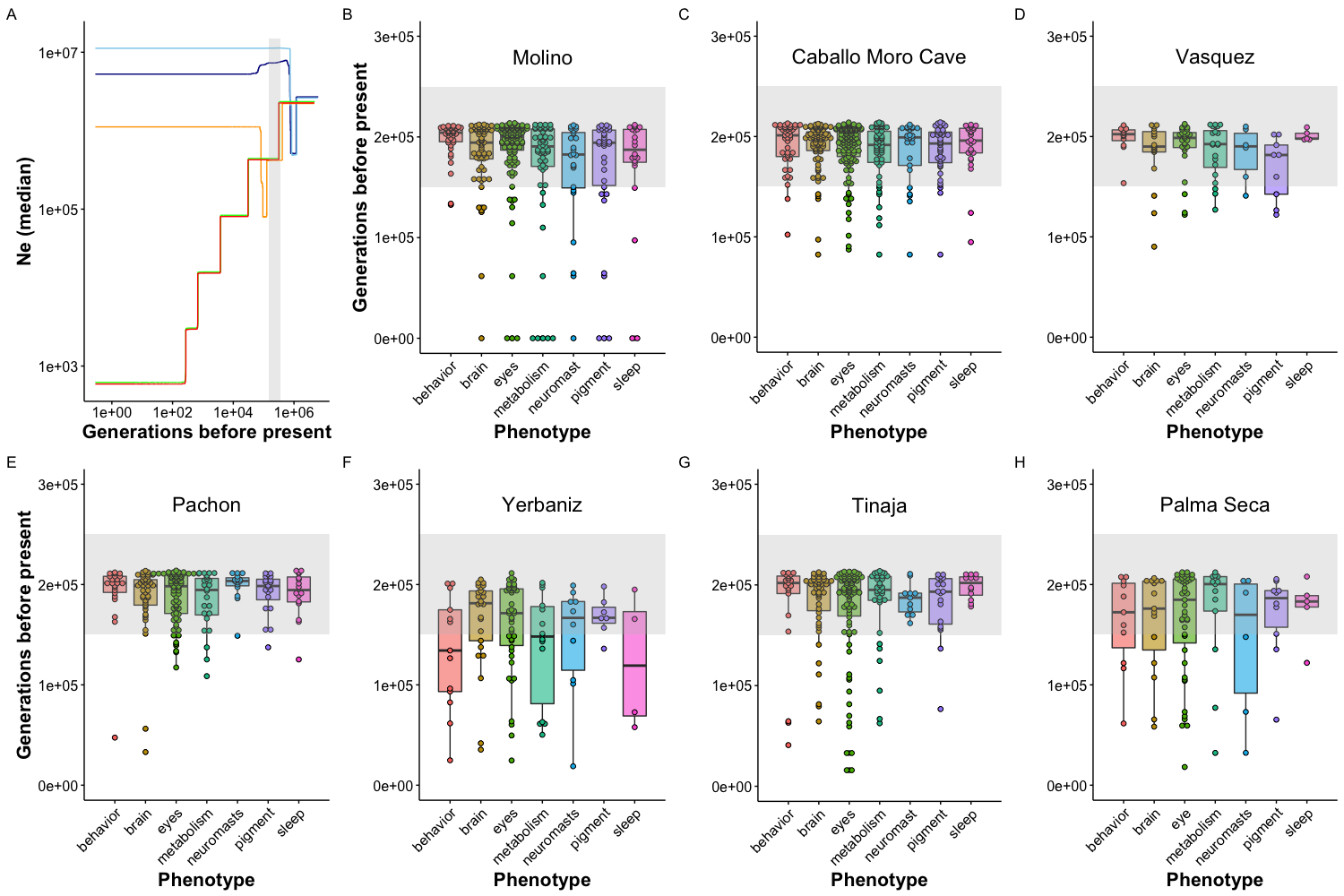

**Rascón**

**Río Choy**

**Pachón**

**Tinaja**

**Molino**

**Figure S12.** Estimated demographic history and ages of selective sweeps for cave populations for expanded set of phenotypes (including behavior and brain). (A) Stairway plot showing median Ne over time in Pachón (orange), Tinaja (red), and Molino (green) cave populations and Rascón (light blue) and Río Choy (dark blue) surface populations. Bottlenecks for present-day cave populations corresponding to the initial cave invasion by ancestral surface stock are highlighted by a gray rectangle (150,000-250,000 generations before present, spanning the range of previous demographic model-based median estimates for split times between Lineage 1 and Lineage 2 cave and surface lineages from (Herman et al. 2018)). Note a more ancient bottleneck in the two surface populations shown (Rascón and Río Choy) around 800,000 generations before present, likely corresponding to migration into northern Mexico. (B-H) Ages of selective sweeps with GO terms associated with cave-adaptive phenotypes (see Table S9) in seven cave populations. Northern lineage caves from the Guatemala regions are shown in B-D. Lineage 2 caves from the El Abra regions are shown in E-H. Gray rectangles span 150,000-250,000 generations before present.

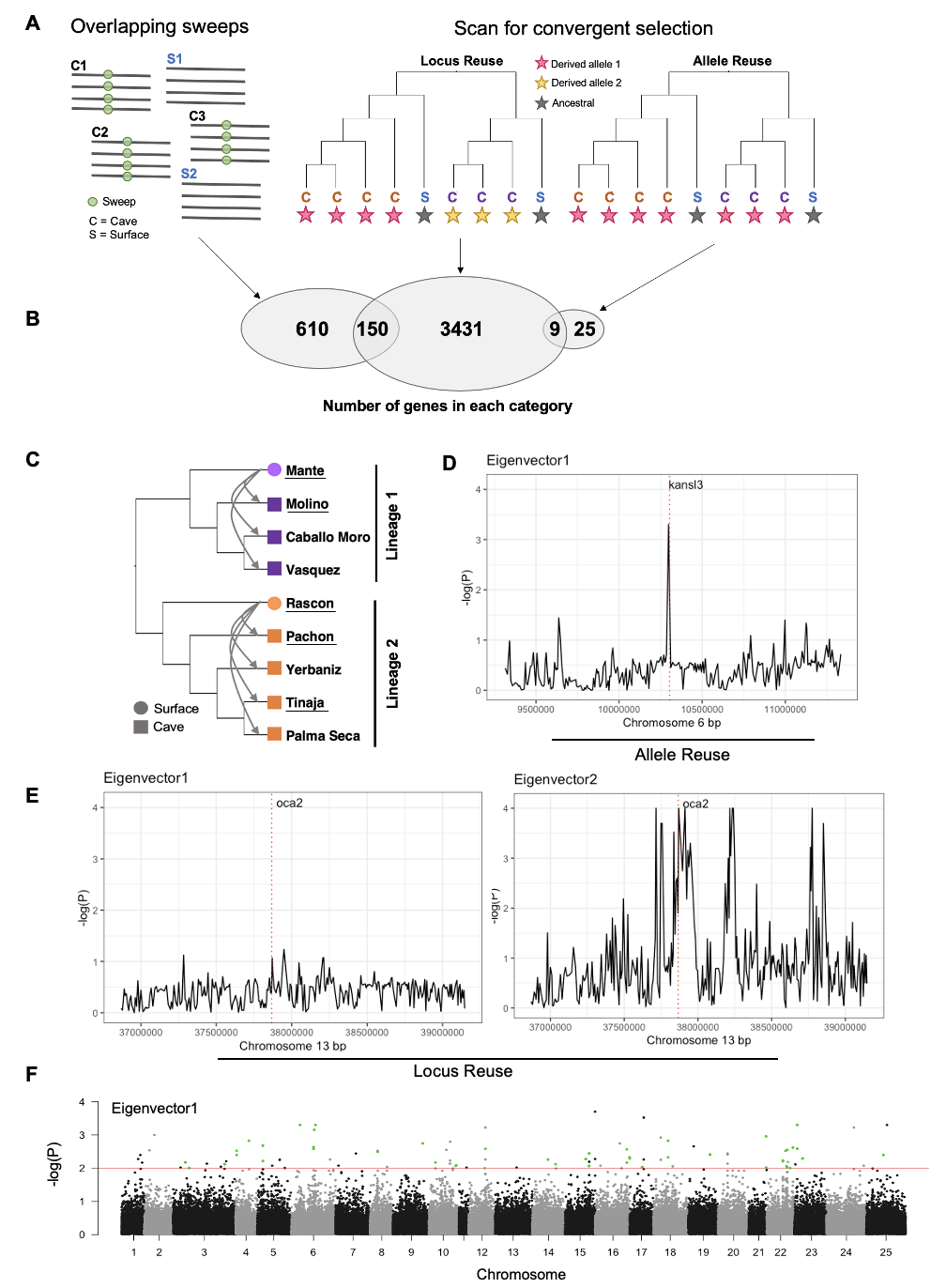

**Figure S13.** Visualization of AFvape-R scan for genomic windows showing a signature of repeated evolution across cavefish lineages. Each point represents a 50 SNP window. Values along the Y axis indicate empirical p-values above the 99th percentile (red line; generated using 10,000 null permutations) for loadings on eigenvector 1 across all 25 chromosomes (see Table S14). The 47 windows (overlapping 34 genes) that showed evidence of allele reuse across all seven cave populations are highlighted in green.

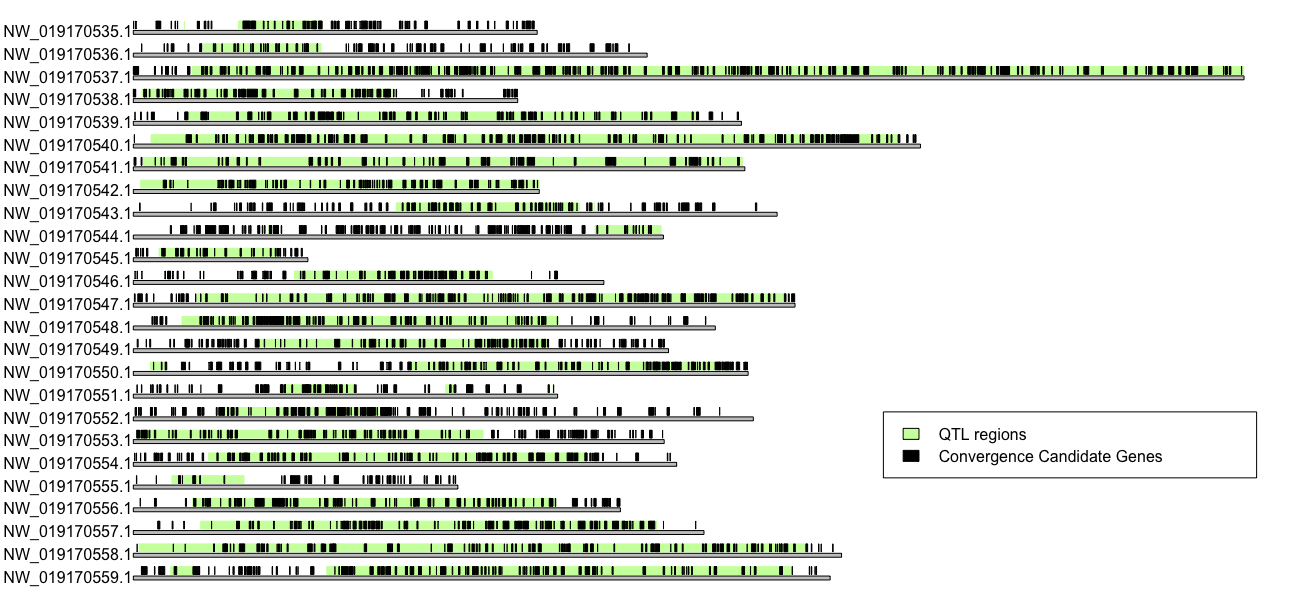

**Figure S14.** Plot of 25 *A. mexicanus* chromosomes with known QTL regions from previous studies highlighted in green and location of 4,085 candidate genes for repeated evolution occurring on an assembled chromosome shown in black. Candidate genes were identified using a scan for parallel selection with AF-vapeR and an overlapping sweeps approach (see main text for details).

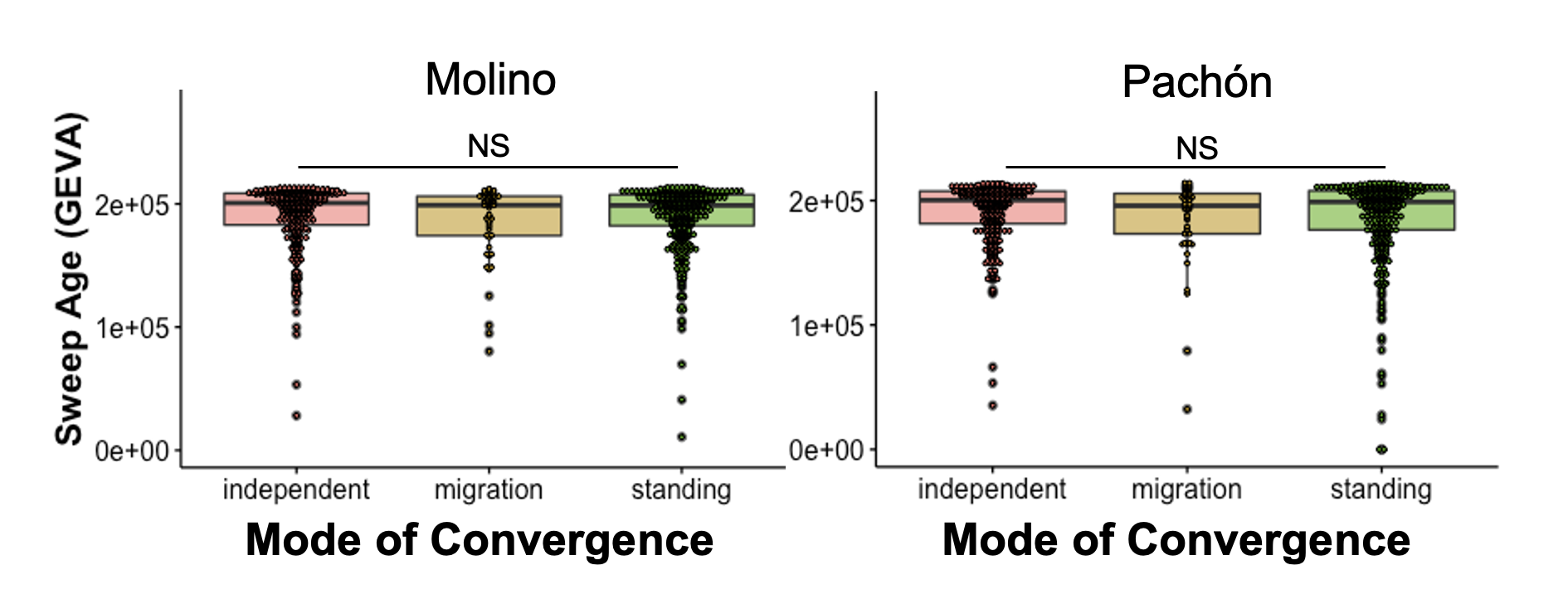
 **Figure S15.** Selective sweep ages (from GEVA) for the 760 overlapping sweep candidate genes do not vary by my mode of repeated evolution (from DMC) in Molino (Lineage 1, left) and Pachón (Lineage 2, right).

Locus reuse candidate genes (AFvape-R scan)

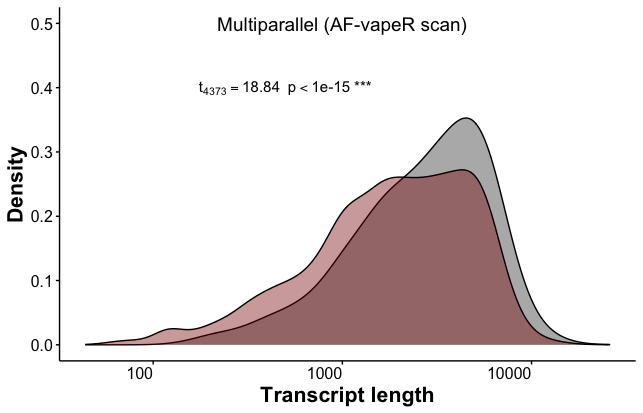

**Figure S16.** Density plot of transcript length for the set candidate genes identified by AF-vapeR as evolving repeatedly via locus reuse across seven cave populations (gray) versus the whole genome (red).

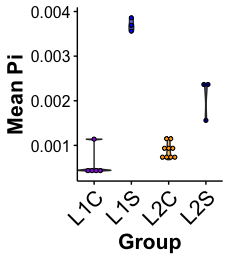

**Figure S17.** Mean nucleotide diversity (pi) within each population, categorized by lineage and ecotype. Each dot represents one population. Hybrid populations (i.e., Caballo Moro Cave eyed individuals, Arroyo, Chica, Toro, and Subterráneo) and populations with less than 3 samples (i.e., Micos and Jalpan) were excluded. L1C = lineage 1 cave, L1S = lineage 1 surface, L2C = lineage 2 cave, L2S = lineage 2 surface.

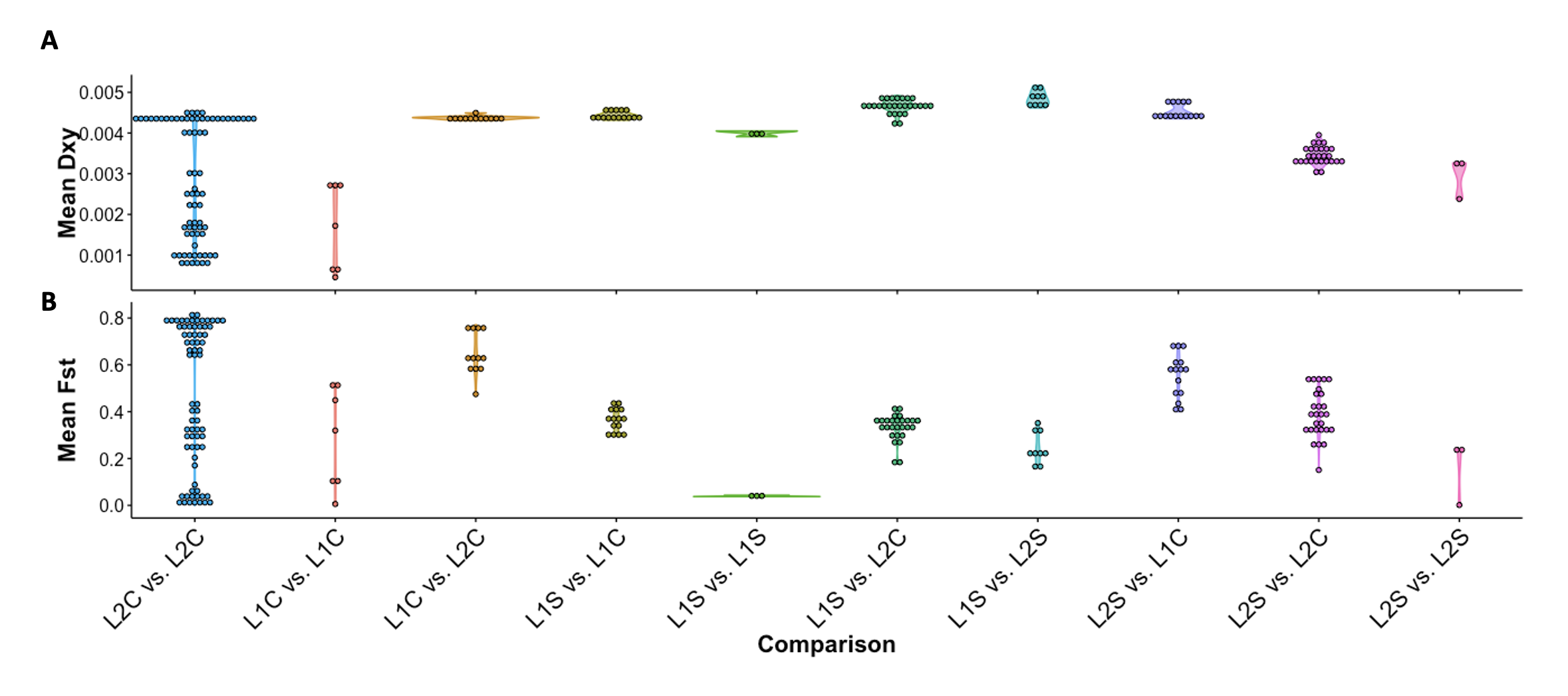

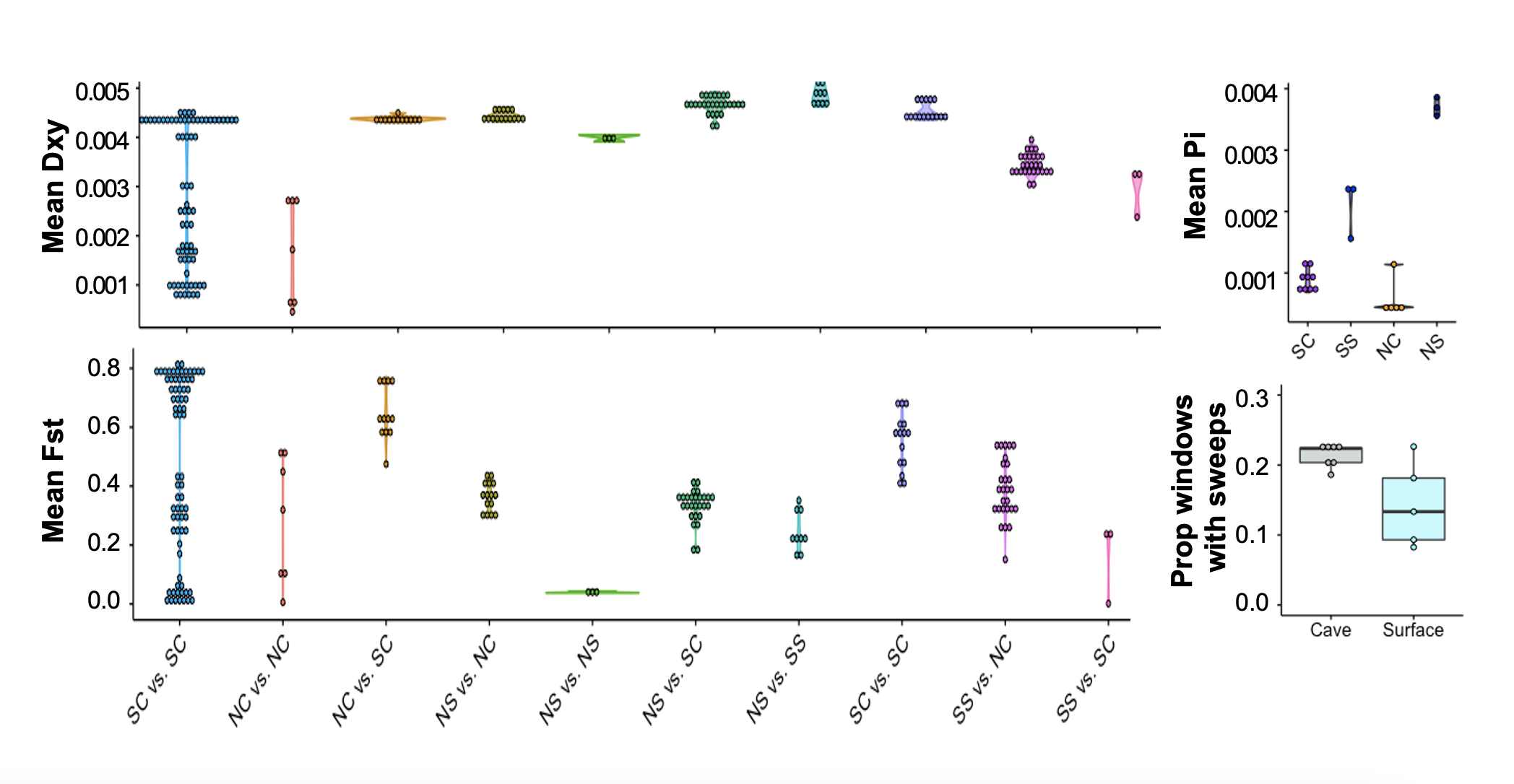

**Figure S18.** (A) Mean Dxy and (B) mean Fst (calculated in 50 kb windows across the genome) between and within ecotypes and lineages. Each dot represents one pairwise population comparison. Hybrid populations (i.e., Caballo Moro Cave eyed individuals, Arroyo, Chica, Toro, and Subterráneo) and populations with less than 3 samples (i.e., Micos and Jalpan) were excluded. L1C = lineage 1 cave, L1S = lineage 1 surface, L2C = lineage 2 cave, L2S = lineage 2 surface.

**Table S1.** Sample IDs, sequencing information, coverage, and raw and clean read counts.

See Excel document.

**Table S2.** Results of D statistic and f4 ratio tests for introgression. *A. nicarageunsis* served as the outgroup. BBAA = derived alleles shared by P1 and P2. ABBA = derived alleles shared by P2 and P3. BABA = derived alleles shared by P1 and P3. Significant p-values (< 0.05) indicate evidence of introgression. An excess of BABA alleles is indicative of introgression between P1 and P2. An excess of ABBA alleles is indicative of introgression between P2 and P3. Subter = Subterranéo. *A. nic = A. nicarageunsis.* Key comparisons between Subterranéo and a new lineage cave (i.e., Escondido) and old lineage caves (i.e., Pachón and Tinaja) highlighted.

| **P1** | **P2** | **P3** | **P4 (outgroup)** | **Dstat** | **Z-score** | **p-value** | **f4-ratio** | **BBAA** | **ABBA** | **BABA** |
| --- | --- | --- | --- | --- | --- | --- | --- | --- | --- | --- |
| Mante | Escondido | Pachón | *A. nic* | 0.1027 | 12.4326 | <0.0001 | 0.0538 | 10,785.00 | 7,257.88 | 5,906.06 |
| Mante | Escondido | Rascón | *A. nic* | 0.2058 | 17.4161 | <0.0001 | 0.1692 | 13,029.60 | 6,028.40 | 3,970.98 |
| Mante | Subter | Escondido | *A. nic* | 0.0655 | 12.2241 | <0.0001 | 0.0308 | 9,921.08 | **8,486.78** | 7,443.59 |
| Mante | Escondido | Tinaja | *A. nic* | 0.1631 | 19.5420 | <0.0001 | 0.0781 | 12,262.50 | 6,474.62 | 4,659.12 |
| Escondido | Pachón | Rascón | *A. nic* | 0.2047 | 16.7233 | <0.0001 | 0.2195 | 7,739.13 | 6,484.09 | 4,280.39 |
| Subter | Escondido | Pachón | *A. nic* | 0.0367 | 5.0787 | <0.0001 | 0.0192 | 11,077.70 | 6,524.09 | 6,062.08 |
| Tinaja | Pachón | Escondido | *A. nic* | 0.2299 | 19.2203 | <0.0001 | 0.0569 | 14,111.20 | 6,011.75 | 3,764.41 |
| Subter | Escondido | Rascón | *A. nic* | 0.0872 | 11.3729 | <0.0001 | 0.0771 | 13,205.80 | 5,215.82 | 4,378.82 |
| Rascón | Tinaja | Escondido | *A. nic* | 0.1241 | 20.0780 | <0.0001 | 0.0294 | 7,072.97 | 5,427.59 | 4,229.47 |
| Subter | Escondido | Tinaja | *A. nic* | 0.0609 | 8.9048 | <0.0001 | 0.0297 | 12,473.50 | 5,662.26 | 5,012.16 |
| Rascón | Pachón | Mante | *A. nic* | 0.3571 | 24.2395 | <0.0001 | 0.2522 | 8,185.19 | 8,099.02 | 3,836.95 |
| Mante | Subter | Pachón | *A. nic* | 0.0718 | 11.2772 | <0.0001 | 0.0350 | 13,325.50 | **6,715.01** | 5,814.88 |
| Tinaja | Pachón | Mante | *A. nic* | 0.2719 | 20.0060 | <0.0001 | 0.1803 | 16,274.10 | 6,511.76 | 3,727.89 |
| Mante | Subter | Rascón | *A. nic* | 0.1294 | 15.1870 | <0.0001 | 0.0992 | 15,954.10 | 5,383.49 | 4,149.91 |
| Rascón | Tinaja | Mante | *A. nic* | 0.1581 | 16.3223 | <0.0001 | 0.0870 | 8,933.86 | 5,400.27 | 3,925.64 |
| Mante | Subter | Tinaja | *A. nic* | 0.1096 | 15.6274 | <0.0001 | 0.0491 | 15,046.40 | **5,916.42** | 4,747.49 |
| Subter | Pachón | Rascón | *A. nic* | 0.2700 | 18.1570 | <0.0001 | 0.2791 | 8,084.42 | 7,270.94 | 4,179.30 |
| Pachón | Tinaja | Rascón | *A. nic* | 0.0860 | 9.7585 | <0.0001 | 0.0800 | 13,023.70 | 4,054.58 | 3,412.69 |
| Tinaja | Pachón | Subter | *A. nic* | 0.2400 | 17.7770 | <0.0001 | 0.0831 | 15,172.70 | 6,439.49 | 3,947.16 |
| Rascón | Tinaja | Subter | *A. nic* | 0.1439 | 17.0269 | <0.0001 | 0.0446 | 7,923.63 | 5,592.76 | 4,185.47 |

**Table S3.** Number and proportion of 5 kb genomic windows with predictions in each category (neutral, linkedSoft, soft sweep, linkedHard, hard sweep) and total windows with evidence of hard or soft sweeps (all sweeps) from diploS/HIC.

| **Lineage** | **Population** | **Type** | **n** | **Total5kbWindowsWithPredictions** | **neutral** | **prop neutral** | **linkedSoft** | **prop linkedSoft** | **soft sweeps** | **prop soft sweeps** | **linkedHard** | **prop linkedHard** | **hard sweeps** | **prop hard sweeps** | **all sweeps** | **prop all sweeps** |
| --- | --- | --- | --- | --- | --- | --- | --- | --- | --- | --- | --- | --- | --- | --- | --- | --- |
| 2 | Rascón | Surface | 13 | 215,196 | 13,979 | 0.0650 | 150,102 | 0.6975 | 10,449 | 0.0486 | 12,074 | 0.0561 | 28,592 | 0.1329 | 39,041 | 0.1814 |
| 2 | Peroles | Surface | 9 | 206,088 | 58,302 | 0.2829 | 95,979 | 0.4657 | 38,567 | 0.1871 | 5,175 | 0.0251 | 8,065 | 0.0391 | 46,632 | 0.2263 |
| 1 | Choy | Surface | 9 | 212,384 | 45,929 | 0.2163 | 137,976 | 0.6497 | 25,319 | 0.1192 | 155 | 0.0007 | 3,005 | 0.0141 | 28,324 | 0.1334 |
| 1 | Mante | Surface | 10 | 216,870 | 14,202 | 0.0655 | 181,146 | 0.8353 | 9,602 | 0.0443 | 1,309 | 0.0060 | 10,611 | 0.0489 | 20,213 | 0.0932 |
| 1 | Caballo Moro | Surface | 6 | 211,924 | 4,281 | 0.0202 | 188,378 | 0.8889 | 4556 | 0.0215 | 1757 | 0.0083 | 12952 | 0.0611 | 17,508 | 0.0826 |
| 2 | Yerbaniz | Cave | 7 | 99,975 | 5,995 | 0.0600 | 38,735 | 0.3874 | 15,874 | 0.1588 | 32,876 | 0.3288 | 6,495 | 0.0650 | 22,369 | 0.2237 |
| 2 | Tinaja | Cave | 17 | 208,538 | 12,165 | 0.0583 | 45,369 | 0.2176 | 18,459 | 0.0885 | 103,452 | 0.4961 | 29,093 | 0.1395 | 47,552 | 0.2280 |
| 2 | Palma Seca | Cave | 8 | 76,410 | 4279 | 0.0560 | 26869 | 0.3516 | 11872 | 0.1554 | 28072 | 0.3674 | 5318 | 0.0696 | 17,190 | 0.2250 |
| 2 | Pachón | Cave | 18 | 200,799 | 7,727 | 0.0385 | 19,919 | 0.0992 | 9,217 | 0.0459 | 132,103 | 0.6579 | 31,833 | 0.1585 | 41,050 | 0.2044 |
| 1 | Vasquez | Cave | 8 | 53,484 | 2,786 | 0.0521 | 10,511 | 0.1965 | 5,061 | 0.0946 | 29,352 | 0.5488 | 5,774 | 0.1080 | 10,835 | 0.2026 |
| 1 | Caballo Moro | Cave | 12 | 189,653 | 7079 | 0.0373 | 35926 | 0.1894 | 15213 | 0.0802 | 104216 | 0.5495 | 27219 | 0.1435 | 42,432 | 0.2237 |
| 1 | Molino | Cave | 15 | 190,397 | 2,030 | 0.0107 | 8,676 | 0.0456 | 3,524 | 0.0185 | 144,262 | 0.7577 | 31,905 | 0.1676 | 35,429 | 0.1861 |

**Table S4.** Detailed information for each gene in the surface fish genome annotation.

See Excel spreadsheet.

Tab 1 –Location of each gene in the genome, gene description, associated GO terms and phenotypes from Ensembl’s Biomart, whether the gene overlaps a previously identified QTL, whether the gene description contains any key words associated with known cave-derived traits, metrics of genetic diversity within (pi) and between (Dxy and Fst) select cave and surface populations from each lineage, and summary of results from selection scans using hapFLK and diploS/HIC.

Tab 2 – SIFT and VEP summary of putatively deleterious mutations for each population.

Tab 3 – 760 overlapping sweeps candidate genes for repeated evolution (evidence of hard or soft sweep from diploS/HIC analysis in Pachón, Tinaja, and Molino; neutral evolution in Mante and Rascón ).

**Table S5.** Breakdown of number of 5 kb windows (out of 267,048 total in the assembly) and genes (out of 26,698 total) included in diploS/HIC analyses for each population. Some 5 kb windows were skipped by diploS/HIC during the prediction step due to missing data or lack of SNPs.

| **Lineage** | **Population** | **Type** | **n** | **Total 5kb Windows with Predictions** | **Prop 5kb Windows with Predictions** | **Genes with Predictions** | **Prop Genes with Predictions** |
| --- | --- | --- | --- | --- | --- | --- | --- |
| 1 | Choy | Surface | 9 | 212,384 | 0.795 | 22,866 | 0.856 |
| 1 | Mante | Surface | 10 | 216,870 | 0.812 | 23,222 | 0.870 |
| 1 | Caballo Moro | Surface | 6 | 211,924 | 0.794 | 23,205 | 0.869 |
| 2 | Peroles | Surface | 9 | 206,088 | 0.772 | 22,390 | 0.839 |
| 2 | Rascón | Surface | 13 | 215,196 | 0.806 | 23,117 | 0.866 |
| 1 | Vasquez | Cave | 8 | 53,484 | 0.200 | 7,741 | 0.290 |
| 1 | Caballo Moro | Cave | 12 | 189,653 | 0.710 | 21,045 | 0.788 |
| 1 | Molino | Cave | 15 | 190,397 | 0.713 | 21,251 | 0.796 |
| 2 | Yerbaniz | Cave | 7 | 99,975 | 0.374 | 13,130 | 0.492 |
| 2 | Tinaja | Cave | 17 | 208,538 | 0.781 | 22,737 | 0.852 |
| 2 | Palma Seca | Cave | 8 | 76,410 | 0.286 | 11,034 | 0.413 |
| 2 | Pachón | Cave | 18 | 200,799 | 0.752 | 22,136 | 0.829 |

**Table S6.** Lists of genes with a sweep in a cave population and neutral evolution in a same-lineage surface population and results of accompanying GO enrichment analyses (using the Gene Ontology Consortium online tool, http://geneontology.org) for individual populations and genes with overlapping sweeps in at least one population from both cavefish lineages.

See Excel spreadsheet.

**Table S7.** List of genes with sweeps in two surface populations (Lineage 1: Mante , Lineage 2: Rascón) and neutral evolution in three cave populations (Lineage 1: Molino, Lineage 2: Pachón and Tinaja), results of GO enrichment analysis (using the Gene Ontology Consortium online tool, http://geneontology.org) for individual populations, and results of DMC (Distinguishing Modes of Convergence) analysis.

See Excel spreadsheet.

**Table S8.** Number of candidate genes for adaptive evolution across seven cave populations (i.e., evidence of a soft or hard selective sweep identified by diploS/HIC in the cave population but no sweep in a same-lineage surface population) and number of those candidate genes with a GO term associated with cave-derived phenotypes.

| **Cave population** | **Lineage** | **Sweep in cave, neutral in same-lineage surface** | **Sweep in cave, neutral in same-lineage surface, and relevant GO term** |
| --- | --- | --- | --- |
| Molino | 1 | 5494 | 285 |
| Vasquez | 1 | 1777 | 82 |
| Caballo Moro | 1 | 6238 | 315 |
| Pachón | 2 | 3903 | 206 |
| Yerbaniz | 2 | 2328 | 100 |
| Tinaja | 2 | 4389 | 217 |
| Palma Seca | 2 | 1984 | 103 |

**Table S9.** Keywords related to known cave-derived phenotypes identified in the GO terms associated with each candidate gene. Candidate genes were defined as a gene with a soft or hard sweep in a given cave population but no sweep in the same-lineage surface population, and a fixed or nearly fixed variant in the cave population. R = regressive traits. C = constructive traits.

| **Phenotypic category** | **Keywords (searched for in GO terms)** |
| --- | --- |
| Eye development (R) | eye, lens, optic, iris, retina, detection of light stimulus, visual, photoreceptor |
| Pigment (R) | pigment, melanin |
| Sleep (R) | sleep, circadian, rhythm, photoperiod, response to light stimulus |
| Metabolism (C) | insulin, glucose, body fat, fat pad, adipose |
| Neuromasts (C) | neuromast, hair cell, lateral line |
| Behavior | behavior |
| Brain | brain, hypothalamus, amygdala, telencephalon |

**Table S10.** GEVA sweep ages for putatively adaptive alleles (i.e., evidence of a selective sweep in a cave population and neutral evolution in a corresponding same-lineage surface population, and a GO term annotation associated with a cave-derived trait category; see Table S9) in each of the seven cave populations analyzed.

See Excel spreadsheet.

**Table S11.** Results of Kruskal-Wallis rank sum tests on estimated timing of selective sweeps (from GEVA) in genes with GO terms associated with cave-derived phenotypic categories (i.e., eyes, pigment, sleep, metabolism, brain, behavior, neuromasts; see Table S10, Figure S11) within each of the seven cave populations examined.

| **Cave Population** | **Chi-square** | **df** | **P** |
| --- | --- | --- | --- |
| Molino | 10.18 | 6 | 0.12 |
| Caballo Moro | 3.18 | 6 | 0.79 |
| Vasquez | 8.47 | 6 | 0.21 |
| Pachón | 3.96 | 6 | 0.68 |
| Yerbaniz | 4.74 | 6 | 0.58 |
| Tinaja | 3.54 | 6 | 0.74 |
| Palma Seca | 2.27 | 6 | 0.89 |

**Table S12.** Results of Wilcoxon rank sum tests on estimated timing of selective sweeps (from GEVA) in genes with GO terms associated with regressive traits (i.e., eyes, pigment, sleep) versus constructive traits (i.e., metabolism, neuromasts; see Table S10, Figure 2B-H) within each of the seven cave populations examined.

| **Cave Population** | **W** | **P** |
| --- | --- | --- |
| Molino | 4996 | 0.94 |
| Caballo Moro | 6423 | 0.77 |
| Vasquez | 530 | 0.52 |
| Pachón | 2060 | 0.44 |
| Yerbaniz | 530 | 0.90 |
| Tinaja | 2487 | 0.50 |
| Palma Seca | 577 | 0.39 |

**Table S13.** DMC results for 794 candidate genes for repeated evolution in caves (760 from overlapping sweeps approach, 150 of which were classified as multiparallel by AF-vapeR; 34 classified as full parallel by AF-vapeR) and for “control” set of 172 genes with evidence of overlapping sweeps in surface populations (Lineage 1: Mante; Lineage 2: Rascón) and neutral evolution in Pachón, Tinaja, and Molino caves. Dxy-based allele split times (between cave and surface alleles) are provided for all 794 candidate genes for repeated evolution in caves. GEVA-based sweep ages are provided for the 760 overlapping sweep candidate genes for repeated evolution in caves.

See Excel spreadsheet.

**Table S14.** AF-vapeR results. The first tab includes all significant windows (above the 99^th^ percentile, P < 0.01). Full parallel windows had significant loadings on Eigenvector 1. Multiparallel windows had significant loadings on Eigenvectors 1 and 2. Divergent (nonparallel) windows had significant loadings on Eigenvectors 1, 2, and 3. Windows with significant antiparallel loadings on any eigenvector were discarded from further analysis. The second and third tabs list genes within significant windows of interest (full parallel and multiparallel, respectively).

See Excel spreadsheet.

**Table S15.** ShinyGO (http://bioinformatics.sdstate.edu/go74/) gene ontology enrichment analysis results for candidate genes for repeated evolution in caves (AF-vapeR allele reuse and locus reuse, and overlapping sweep candidate genes) and for genes showing a pattern of repeated evolution in surface populations.

See Excel spreadsheet.

**Table S16.** Effect sizes (Cohen's *d*) for comparison of transcript length in candidate genes for repeated evolution in caves, broken down by mode of repeated evolution predicted with DMC, compared to the whole genome.

| **Predicted Mode of Repeated Evolution** | **Cohen's *d*** |
| --- | --- |
| Independent | 0.524 |
| Migration | 0.264 |
| Standing | 0.256 |

**Table S17.** GATK filters applied to variant and invariant sites. QD = QualByDepth. FS = FisherStrand. MQ = RMSMappingQuality.

| **Invariant sites** | **SNPs** | **Mixed/indels** |
| --- | --- | --- |
| QD < 2.0 | QD < 2.0 | QD < 2.0 |
| FS > 60.0 | FS > 200.0 | FS > 200.0 |
| MQ < 40.0 | ReadPosRankSum < -20.0 | ReadPosRankSum < -20.0 |

**Table S18.** Information on the number of invariant sites and variant sites (including SNPs, indels, and mixed sites) per chr.

See Excel document.

**Table S19.** Mean, median, min, and max Pi within each population and Fst and Dxy between each pair of populations calculated in 50 kb windows across the genome.

See Excel document.

**Table S20.** Parameter values specified to DMC. sels = selection coefficient; times = standing time; gs = frequency of the standing variant; migs = migration rate; proportion of migrants between populations each generation

| **DMC Parameter** | **Values Specified** |
| --- | --- |
| sels | 0.0001, 0.0010, 0.0100, 0.0200, 0.0300, 0.0400, 0.0500, 0.0600, 0.0700, 0.0800, 0.0900  0.1000, 0.1100, 0.1200, 0.1300, 0.1400, 0.1500, 0.2000, 0.2500, 0.3000, 0.4000, 0.5000,  0.6000 |
| times | 5, 50, 100, 500, 1000, 10000, 1000000 |
| gs | 0.00005, 0.0001, 0.001, 0.01, 0.1 |
| migs | 0.00001, 0.001, 0.1, 0.5, 1.0 |
